# Mechanosensitive Stanniocalcin-1 Attenuates Pulmonary Arterial Hypertension by Suppressing Smooth Muscle Cell Proliferation

**DOI:** 10.1101/2025.10.13.682225

**Authors:** Mariko Kogami, Yuko Kato, Satoko Ito, Keiko Uchida, Hana Inoue, Shota Tanifuji, Mayumi Yokotsuka, Yoshinari Yamamoto, Toshitaka Nagao, Shinji Abe, Roger R. Reddel, Kazufumi Nakamura, Utako Yokoyama

## Abstract

**Background:** Idiopathic pulmonary arterial hypertension (IPAH) is driven by progressive pulmonary vascular remodeling, particularly pulmonary arterial smooth muscle cell (PASMC) proliferation. Current combination vasodilator therapies have markedly improved outcomes; however, prognosis remains poor in subgroups such as patients with respiratory comorbidities, highlighting the need for novel therapies. Elevation of intravascular hydrostatic pressure is a hallmark of IPAH, yet its direct role in PASMC pathobiology remains largely unexplored. We developed a cyclic hydrostatic pressurization culture system to model hypertensive hemodynamic stress *in vitro* and identify pressure-responsive mediators.

**Methods:** PASMCs from 4 patients with IPAH were exposed to high hydrostatic pressure (70/40 mmHg, 60 bpm). Transcriptomic profiling identified differentially expressed genes, validated by qPCR. Functional studies included piezo-type mechanosensitive ion channel component 1 (PIEZO1) modulation, recombinant human stanniocalcin-1 (rhSTC1) treatment, bromodeoxyuridine (BrdU) incorporation, and western blotting for cell-cycle regulators. *In vivo*, chronic hypoxia–induced pulmonary hypertension was assessed in wild-type and *Stc1*^−/−^ mice by hemodynamic and histological analyses, with or without intratracheal rhSTC1 administration.

**Results:** RNA sequencing revealed *STC1* as a robustly pressure-induced gene in IPAH PASMCs. PIEZO1 activation upregulated *STC1*, whereas knockdown blunted this response. Elevated *STC1* expression was observed in PASMCs of IPAH lung tissues, and rhSTC1 reduced PASMC proliferation and increased p-p53, p21, and p27 expression. In the chronic hypoxia model, *Stc1*^−/−^ mice exhibited higher right ventricular systolic pressure (RVSP) (43.7 ± 1.3 vs. 30.6 ± 0.9 mmHg) and greater pulmonary arterial medial thickness (39.1 ± 2.5% vs. 26.5 ± 1.3%) than wild-type mice. CD68-positive macrophages were increased in *Stc1*^−/−^ mice under normoxia and further elevated with hypoxia. In wild-type and *Stc1*^−/−^ PAH models, intratracheal administration of rhSTC1 markedly reduced medial thickening, CD68-positive macrophage accumulation, and RVSP in both wild-type and *Stc1*^−/−^ mice.

**Conclusions:** We demonstrate that elevated hydrostatic pressure drives *STC1* expression via PIEZO1, conferring potent anti-remodeling effects in IPAH. STC1 supplementation represents a potential therapeutic strategy that addresses an urgent medical need not fulfilled by conventional therapies.

**Clinical Perspective:** *What Is New?:*

- We established a novel hydrostatic pressurization system to recapitulate idiopathic pulmonary arterial hypertension (IPAH) hemodynamic conditions *in vitro*.
- Stanniocalcin-1 (*STC1*) is a hydrostatic pressure–responsive gene in pulmonary arterial smooth muscle cells (PASMCs) from IPAH patients, induced via the mechanosensitive receptor piezo type mechanosensitive ion channel component 1 (PIEZO1).
- Exogenous STC1 suppresses PASMC proliferation and attenuates pulmonary vascular remodeling in chronic hypoxia–induced PAH models.

*What Are the Clinical Implications?:*

- STC1 supplementation represents a potential therapeutic strategy for IPAH, acting through a non-vasodilatory mechanism.
- STC1 supplementation may offer benefit in patients with limited response to current vasodilator therapies or with comorbid respiratory disease.
- Targeting mechanotransduction pathways could expand treatment options for pulmonary hypertension.

## Introduction

Pulmonary arterial hypertension (PAH) is a progressive and life-threatening disease characterized by elevated pulmonary arterial pressure, right ventricular hypertrophy, and extensive pulmonary vascular remodeling.^1^ Among the various cell types implicated in PAH pathogenesis, pulmonary arterial smooth muscle cells (PASMCs) play an essential role by undergoing phenotypic modulation and hyperproliferation, ultimately contributing to vascular occlusion and increased pulmonary vascular resistance.^2^ Although vasodilator therapies have advanced, current treatments largely target vasoconstriction and endothelial dysfunction. Efforts to develop therapies that address structural vascular remodeling are ongoing; however, overcoming this aspect of the disease remains a significant challenge.

Mechanical stress has emerged as a critical regulator of vascular cell behavior and remodeling in cardiovascular diseases.^3^ Although the effects of cyclic stretch and shear stress have been extensively studied in the pulmonary circulation,^3^ the contribution of intraluminal hydrostatic pressure—defined as the force exerted perpendicularly on the vessel wall—remains poorly understood. Hydrostatic pressure has been shown to influence endothelial cell morphology, proliferation, calcium signaling, extracellular matrix composition, adhesion molecule expression, and barrier permeability,^4^ all of which are implicated in endothelial dysfunction—a common feature of hypertension, atherosclerosis, stroke, and pathological angiogenesis. However, despite its direct impact on vascular wall, the effect of hydrostatic pressure on vascular smooth muscle cells (VSMCs)—the most abundant structural component of the vascular wall—has received little attention. Notably, VSMCs are highly responsive to their local microenvironment, integrating mechanical cues, soluble factors, cell–matrix interactions, and inflammatory mediators.^5^ They exhibit remarkable phenotypic plasticity, shifting from a quiescent, contractile state to a synthetic, proliferative phenotype under pathological conditions.^6^ Based on these characteristics, we hypothesized that VSMCs may act as central mediators of hydrostatic pressure–induced remodeling in PAH.

We previously developed a custom-built culture system capable of applying cyclic hydrostatic pressure to cells.^7,8^ For the present study, we modified the system to reproduce both the magnitude and frequency of cyclic hydrostatic pressure characteristic of PAH, thereby enabling more physiologically relevant modeling of the disease condition. Using this refined system, we exposed PASMCs derived from patients with idiopathic pulmonary arterial hypertension (IPAH) to pressure cycles that mimic hypertensive hemodynamic stress. Subsequently, transcriptomic profiling was performed to identify pressure-sensitive genes selectively upregulated in IPAH PASMCs. Functional analyses were conducted to determine the roles of these genes in driving vascular remodeling. Our findings uncovered a previously unrecognized pressure-responsive mechanotransduction pathway in human PASMCs.

## Methods

The full experimental details are presented in the Supplemental Materials.

### Study Design

This preclinical study used patient-derived PASMCs and mice models of pulmonary hypertension. *In vitro*, gene expression changes under mechanical stimulation were assessed using a custom-built hydrostatic pressure apparatus. Pressure-responsive molecules identified *in vitro* were further evaluated *in vivo* using PASMCs from patients with IPAH, a hypoxia-induced pulmonary hypertension mouse model, and genetically modified mice.

### Human Samples

PASMCs and lung tissues from IPAH patients and non-PAH controls were obtained at Okayama University with institutional review board approval and written informed consent (reference number: 1511-017). Donor characteristics are provided in Table S1. In addition, commercially available non-PAH control cells were used, including PASMCs from a 56-year-old female donor (non-PAH-8, CC-2581; Lonza).

### Hydrostatic Pressure Stimulation

The hydrostatic pressure apparatus consisted of a pressure chamber, regulator, and compressor (Figure S1A-C). Cell culture dishes were placed inside the pressure chamber, and air drawn from a CO_2_ incubator was periodically introduced into the chamber via the regulator to apply cyclic hydrostatic pressure to the cells. PASMCs were seeded at 1.5 × 10^5^ cells per 35-mm dish and placed in the pressure chamber. Cyclic hydrostatic pressure of 70/40 mmHg (mean 50 mmHg) at 60 bpm was applied for 72 h. Pressure waveforms were verified using a pressure transducer (Figure S1B). Control cells were cultured under atmospheric pressure in the same CO_2_ incubator.

### RNA Sequencing

Total RNA was extracted from cultured PASMCs using the RNeasy Mini Kit (Qiagen) according to the manufacturer’s instructions, and RNA integrity was confirmed (RIN > 7.0). Library preparation, sequencing, and primary processing were performed by Rhelixa (Tokyo, Japan). Poly(A)-selected libraries were sequenced on an Illumina NovaSeq X Plus platform to generate paired-end 150-bp reads at a depth of approximately 26.7 million reads/sample. Reads were quality-checked, adapter-trimmed, and aligned to the GRCh38 reference genome. Differential expression analysis was performed using DESeq2, and gene set enrichment analysis using fgsea. Transcription factor enrichment analysis was performed with iRegulon.

### Mouse Models

Wild-type (WT) and Stanniocalcin-1(*Stc1)*⁻^/^⁻ mice on a C57BL/6J background (male and female, 3 months old)^9^ were maintained under normoxic (21% O_2_) or hypoxic (10% O_2_) conditions for 4 weeks. For therapeutic studies, recombinant human stanniocalcin-1 (rhSTC1) was administered intratracheally (2 µg/50 µL PBS, every 72 h for 4 weeks) based on previous reports^10^ and pharmacokinetic pilot data (Figure S2).

### Hemodynamics and Morphometry

Echocardiography and left/right heart catheterization were performed to assess hemodynamics. Fulton index (right ventricle/[left ventricle + septum]; RV/[LV+S]) was calculated from heart weights. Paraffin-embedded lung sections were stained with Elastica–Masson–Goldner to quantify medial wall thickness: ([external elastic lamina diameter – internal elastic lamina diameter]/external elastic lamina diameter) × 100%. Medial wall thickness was calculated for each pulmonary artery, and the average of 30 vessels with a diameter <100 µm was taken as the representative value for each mouse.

### Molecular Assays

qPCR (SYBR), western blot, ELISA, BrdU incorporation assay, immunohistochemistry, immunofluorescence, and terminal deoxynucleotidyl transferase dUTP nick end labeling (TUNEL) staining were performed as described previously.^11–13^ Primer/probe sequences and antibody details are provided in Tables S2 and S3.

### Statistics

Analyses were performed using GraphPad Prism (v9.2.0). Two-group comparisons were performed using the Mann–Whitney U test, whereas multiple-group comparisons were assessed using the Kruskal–Wallis test followed by post hoc Mann–Whitney U tests. Data are expressed as mean ± SEM. *P*<0.05 was considered statistically significant. Significance is indicated as \**P*<0.05, \*\**P*<0.01, and \*\*\**P*<0.001.

### Study Approval

Human studies were approved by the Ethical Guidelines Committee of Tokyo Medical University and Okayama University (T2023-0084 and 1511-017, respectively) and adhered to the principles outlined in the Declaration of Helsinki (10th version, Fortaleza, 2013). Animal studies were approved by the Tokyo Medical University Institutional Animal Care and Use Committee (R7-058) and were conducted in accordance with the Animal Research: Reporting of In Vivo Experiments (ARRIVE) guidelines.

## Results

### Cyclic Hydrostatic Pressurization Induces Expression of Genes Associated with Proliferation and Dedifferentiation in IPAH-Derived PASMCs

To examine how hydrostatic pressure affects gene expression, PASMCs derived from patients with IPAH were subjected to cyclic hydrostatic pressurization (70/40 mmHg, 60 bpm) for 72 h, followed by RNA sequencing (Figure 1A).

**Figure 1.**
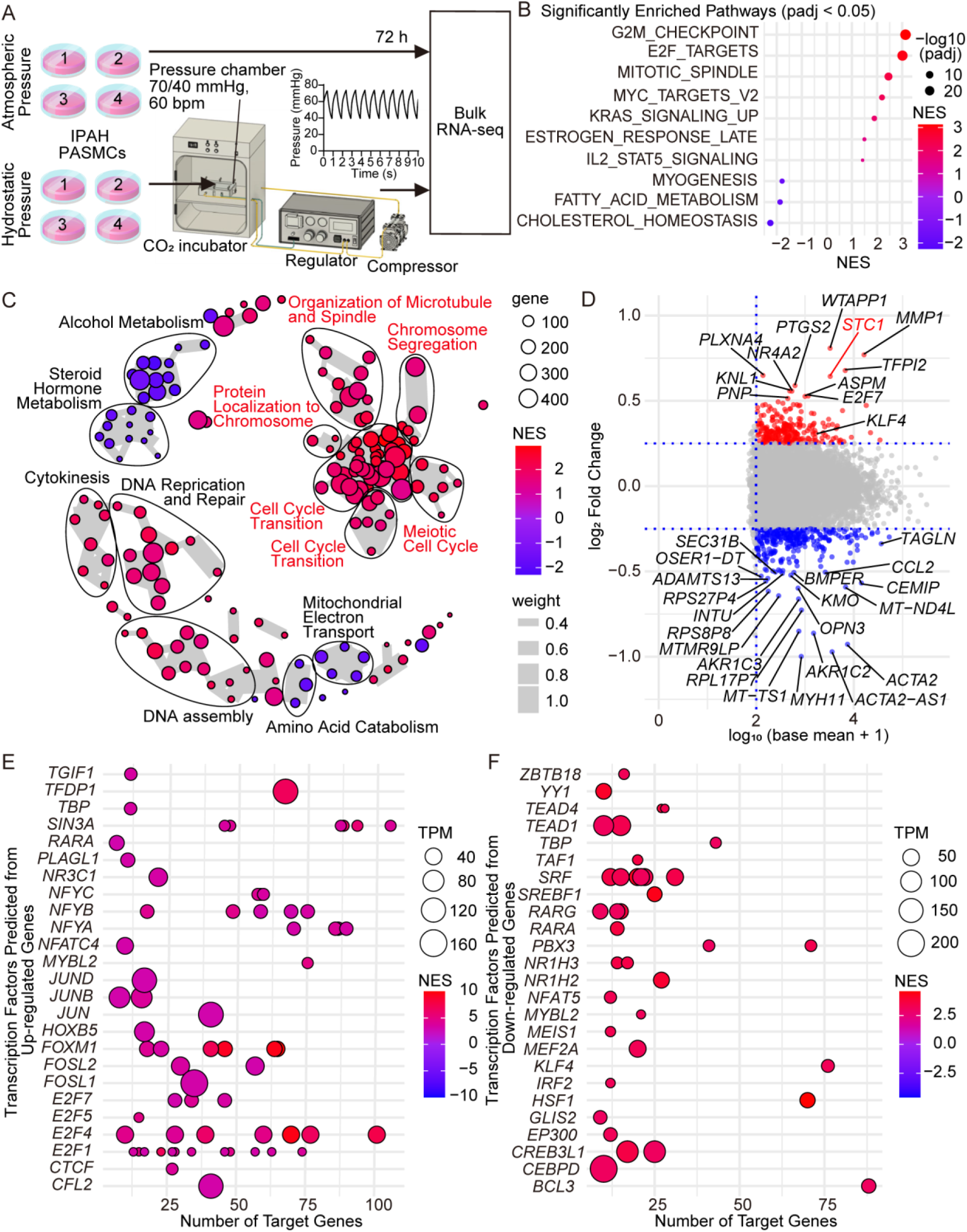
Transcriptomic changes in pulmonary artery smooth muscle cells (PASMCs) subjected to cyclic hydrostatic pressure. **A**, Experimental protocol for cyclic hydrostatic pressure loading. PASMCs derived from patients with idiopathic pulmonary arterial hypertension (IPAH; n=4) were cultured to approximately 80% confluence and then subjected to cyclic hydrostatic pressure (70/40 mmHg , 60 bpm) for 72 h using a custom-designed system. Cells were subsequently harvested for RNA sequencing (RNA-seq). **B**, Dot plots show the top 10 enriched gene sets with adjusted *P*-value (padj) < 0.05 for Kyoto Encyclopedia of Genes and Genomes (KEGG). Circle size represents the number of genes in each gene set, and normalized enrichment score (NES) indicates the degree of enrichment. **C**, Enrichment map generated from Gene Ontology Biological Process (GOBP) gene set enrichment analysis (GSEA) comparing hydrostatic pressure (HP) and atmospheric pressure (AP). Clusters with ≥3 nodes were annotated. **D**, MA plot of RNA-seq data, showing log_2_ fold change versus log_10_(base mean + 1) expression levels. Genes with log₂ fold change >0.5 (red) or <–0.5 (blue) and log_10_(base mean + 1) ≥2 are highlighted. **E**, **F**, Dot plots generated by iRegulon analysis. **E**, Transcription factors (TFs) predicted from upregulated genes; **F**, TFs predicted from downregulated genes.

Gene set enrichment analysis (GSEA) using Kyoto Encyclopedia of Genes and Genomes (KEGG) databases revealed significant enrichment of proliferation-related pathways such as “G2M checkpoint” and “E2F targets” under cyclic hydrostatic pressure (normalized enrichment score [NES] > 2.0, Figure 1B). To gain a comprehensive understanding of the biological processes involved, we conducted Gene Ontology Biological Process (GOBP) enrichment analysis. Consistently, the GOBP enrichment network map revealed that clusters of cell cycle–related gene sets were predominantly upregulated, which are highlighted in red for visualization (Figure 1C).

Mean–average (MA) plot analysis identified 250 upregulated and 291 downregulated genes with log_2_ fold change > 0.25 or < –0.25 and log_10_(base mean + 1) > 2.0 (Figure 1D, Tables S4 and S5). Expression of the key contractile phenotype markers myosin heavy chain 11 (*MYH11*), smooth muscle α-actin (*ACTA2*), and transgelin (*TAGLN*) was consistently downregulated. Concomitantly, Krüppel-like factor 4 (*KLF4*), a transcription factor recognized as a canonical marker of dedifferentiation,^14^ was upregulated. Transcription factor analysis using iRegulon identified transcription factors predicted from upregulated genes (Figure 1E) as well as those associated with downregulated genes (Figure 1F). FOS like 1 (*FOSL1*), a component of the activator protein-1 (AP-1) complex known to promote cell proliferation and phenotypic dedifferentiation,^15^ was predicted as a regulator of upregulated genes. *KLF4* was predicted as a regulator of downregulated genes. These data suggest that elevating intravascular pressure directly promotes a phenotypic shift toward dedifferentiation in PASMCs. To explore a regulator for pressure-induced cell proliferation, we focused on the glycoprotein STC1, which was strongly induced and has been implicated in paracrine/autocrine regulation of cell proliferation.^16–17^

### STC1 Expression Is Elevated in IPAH Patients and Hypoxia-Induced Pulmonary Hypertension Model Mice

Immunohistochemistry of human lung tissues demonstrated markedly increased expression of STC1 in pulmonary arteries of IPAH patients compared with non-PAH controls (Figure 2A). Consistently, western blot analysis of culture supernatants showed significantly higher levels of secreted STC1 in PASMCs from IPAH patients than in those from non-PAH controls (Figures 2B-C). Basal mRNA expression analysis also demonstrated elevated *STC1* levels in IPAH-derived PASMCs compared with non-PAH PASMCs under unstimulated conditions (Figure 2D). Exposure to cyclic hydrostatic pressure for 72 h did not change *STC1* expression levels in the non-PAH PASMCs (Figure 2E); however, PASMCs from IPAH patients exhibited a significant pressure-induced upregulation of *STC1* mRNA (Figures 2F-J).

**Figure 2.**
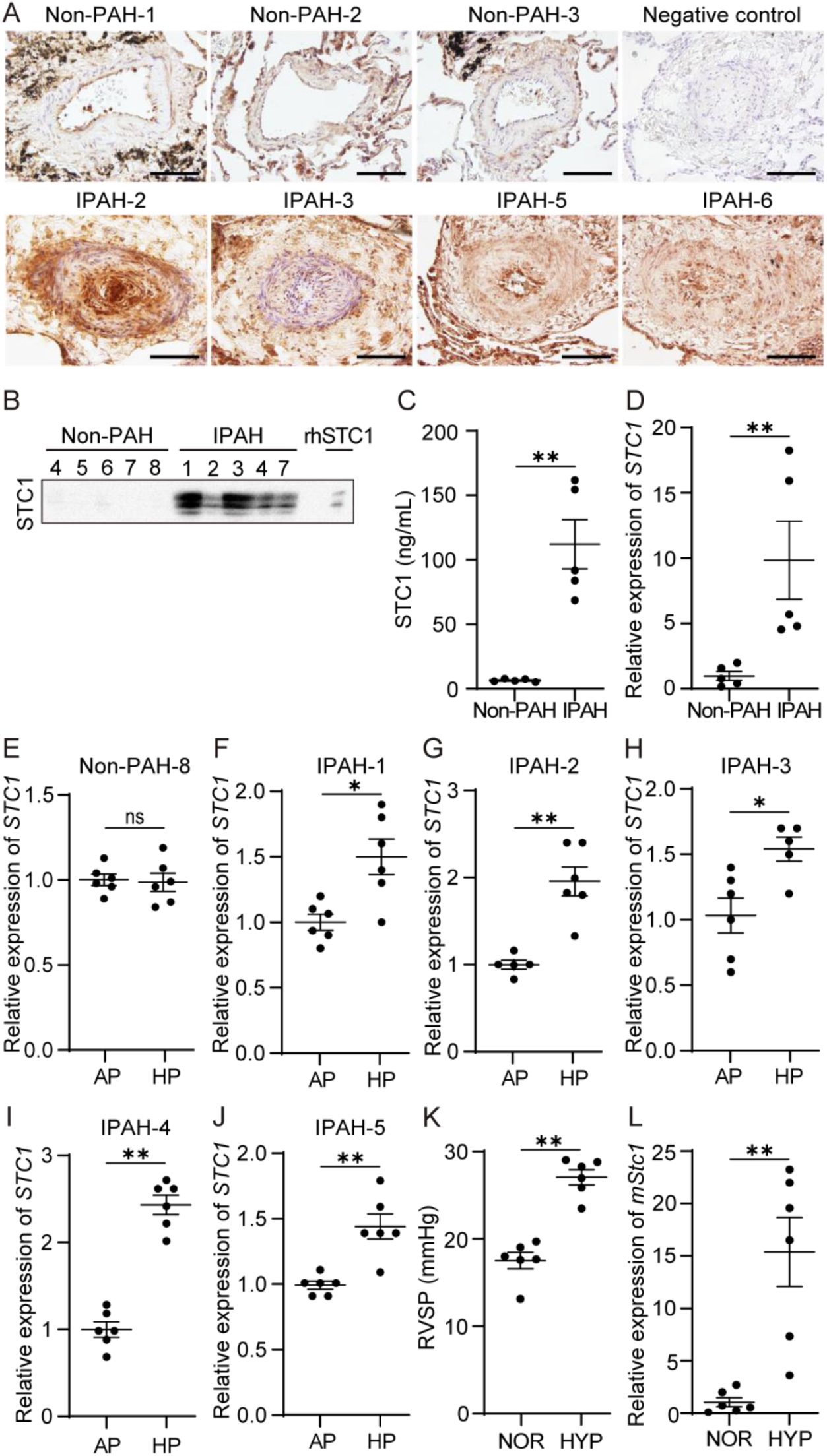
Expression of STC1 in PASMCs from IPAH patients and lungs from hypoxia-induced PAH model mice. **A**, Immunohistochemistry of stanniocalcin-1 (STC1) expression in pulmonary arteries from donors (n=3) and patients with idiopathic pulmonary arterial hypertension (IPAH; n=4). Scale bars: 50 μm. **B**-**C**, Western blot analysis of STC1 in the culture supernatant of pulmonary artery smooth muscle cells (PASMCs) from non-PAH controls (n=5) and IPAH patients (n=5). PASMCs were grown in serum-free Dulbecco’s modified Eagle medium (DMEM) for 24 h. The supernatant was collected and concentrated 22-fold using a column concentrator. Representative blot (**B**) and quantification of STC1 concentrations based on band density measurements using reference bands generated from recombinant human STC1 (rhSTC1) at 500 µg/mL (**C**). **D**, Basal expression of *STC1* mRNA in PASMCs under unstimulated conditions. PASMCs derived from non-PAH controls (n=5) and IPAH patients (n=5) were analyzed by quantitative polymerase chain reaction (qPCR). **E**-**J**, Relative expression of *STC1* mRNA in PASMCs derived from non-PAH control (**E**) and IPAH patients (**F-J**) after exposure to cyclic hydrostatic pressure for 72 h. AP, atmospheric pressure; HP, hydrostatic pressure. Right ventricular systolic pressure (RVSP, **K**), and relative expression of *Stc1* mRNA in lung (**L**) in hypoxia-induced PAH model mice (C57BL/6J; fraction of inspired oxygen [FiO_2_] 10% for 4 weeks; n=6; NOR, normoxia; HYP, hypoxia). Data are presented as mean ± SEM. Statistical significance was determined using the Mann–Whitney U test. \**P* < 0.05, \*\**P* < 0.01, ns: not significant.

Similarly, in a mouse model of hypoxia-induced pulmonary hypertension (PH), characterized by significantly elevated right ventricular systolic pressure (RVSP, Figure 2K), pulmonary *Stc1* mRNA levels were markedly increased (Figure 2L).

### Mechanotransduction via PIEZO1 and HIF-1α Signaling Regulates STC1 Expression

To investigate the mechanism by which hydrostatic pressure induces *STC1* expression, IPAH-derived PASMCs were stimulated with agonists for mechanosensitive channels: transient receptor potential vanilloid 4 (TRPV4; GSK1016790A), transient receptor potential melastatin 7 (TRPM7; naltriben), and PIEZO1 (Yoda1). Among these, PIEZO1 activation significantly increased *STC1* expression in PASMCs from 2 independent IPAH patients (Figure 3A). Small interfering RNA (siRNA)–mediated knockdown of *PIEZO1* effectively suppressed the hydrostatic pressure–induced upregulation of *STC1* (Figures 3B-C). Immunohistochemistry of human lung tissues demonstrated that PIEZO1 expression was increased in IPAH patients compared with non-PAH controls, although control samples exhibited some variability in expression levels (Figure 3D). These findings are consistent with a previous report.^18^

**Figure 3.**
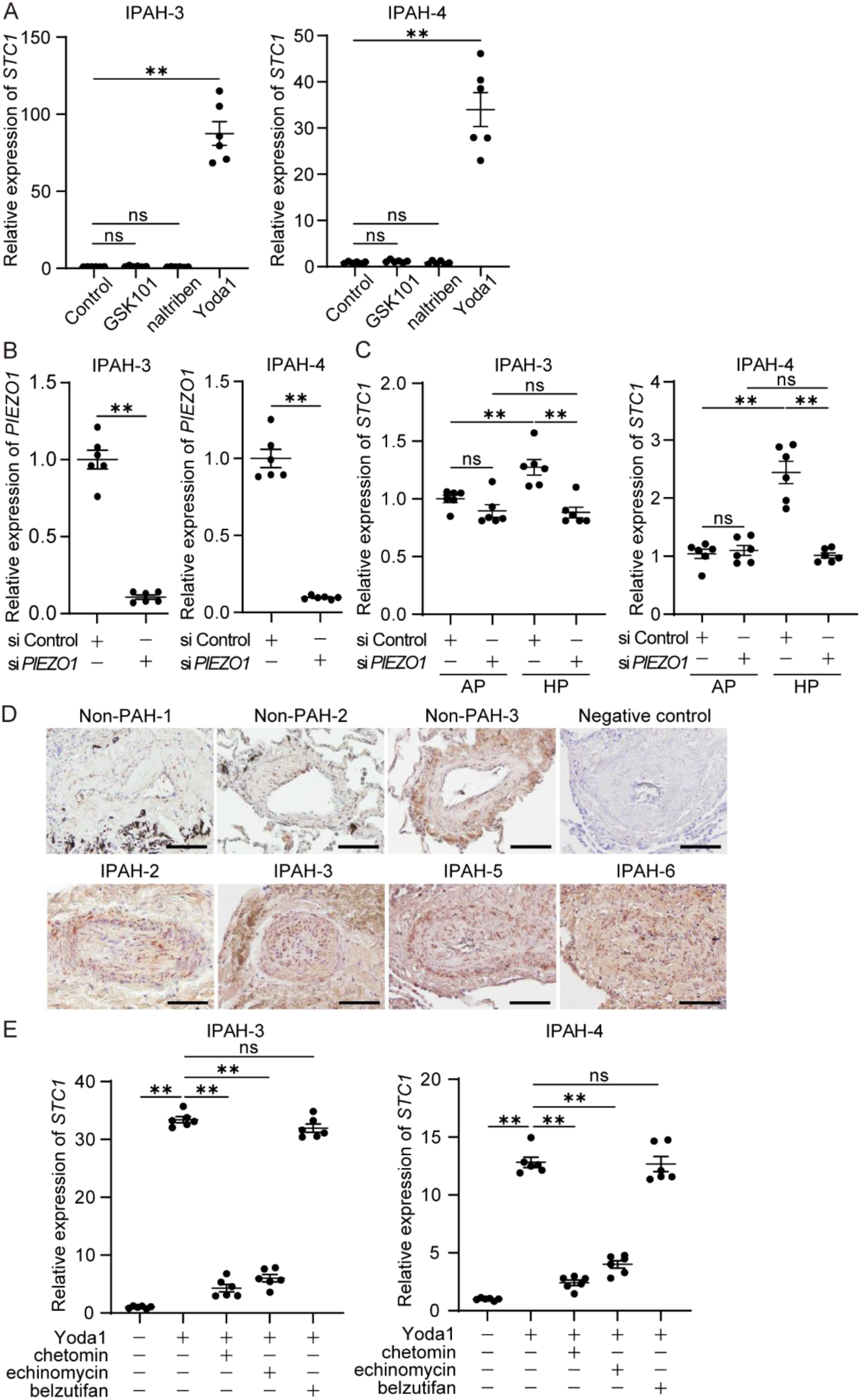
Regulation of STC1 expression by mechanotransduction. **A**, Effects of induction of *STC1* expression by mechanoreceptor activators. Relative expression of *STC1* mRNA following stimulation of mechanosensitive receptors in pulmonary artery smooth muscle cells (PASMCs) from 2 patients with idiopathic pulmonary arterial hypertension (IPAH; left, IPAH-3; right, IPAH-4). Cells were stimulated with GSK1016790A (10 nmol/L), an agonist of transient receptor potential vanilloid 4 (TRPV4); naltriben (20 μmol/L), an agonist of transient receptor potential melastatin 7 (TRPM7); or Yoda1 (20 μmol/L), an agonist of PIEZO1. *STC1* mRNA expression was quantified using quantitative polymerase chain reaction (qPCR). **B**, Efficiency of *PIEZO1* knockdown. PASMCs were transfected with small interfering RNA (siRNA) targeting *PIEZO1* (si *PIEZO1*) or control siRNA (si Control), and knockdown efficiency was assessed by qPCR. Relative expression of *PIEZO1* mRNA in IPAH-3 (left) and IPAH-4 (right). **C**, Effect of *PIEZO1* knockdown on *STC1* expression under cyclic hydrostatic pressure. Relative expression of *STC1* mRNA in PASMCs from IPAH patients (left, IPAH-3; right, IPAH-4) following exposure to cyclic hydrostatic pressure for 72 h with or without *PIEZO1* knockdown. AP, atmospheric pressure; HP, hydrostatic pressure. AP**D**, Immunohistochemistry of PIEZO1 expression in pulmonary arteries from non-PAH controls (n=3) and patients with idiopathic IPAH (n=4). Scale bars: 50 μm. **E**, Effect of hypoxia-inducible factor (HIF) inhibition on PIEZO1-mediated STC1 upregulation. Relative expression of *STC1* mRNA in PASMCs from IPAH patients (left, IPAH-3; right, IPAH-4) after treatment with HIF inhibitors during PIEZO1-mediated upregulation of STC1. HIF-1α was inhibited using chetomin (100 nmol/L) or echinomycin (30 nmol/L), and HIF-2α was inhibited using belzutifan (10 nmol/L). PIEZO1 activation was treated by Yoda1. Expression levels were quantified by qPCR. Data are presented as mean ± SEM (n=6). For comparisons among multiple groups, statistical significance was assessed using the Kruskal-Wallis test followed by post hoc Mann–Whitney U tests. Comparisons between two groups were performed using the Mann–Whitney U test. \*\**P* < 0.01, ns: not significant.

Previous studies demonstrated that PIEZO1 activation increases hypoxia-inducible factor-1α (HIF-1α) expression.^19–21^ To investigate whether PIEZO1-mediated induction of *STC1* depends on HIF signaling, cells were treated with chetomin and echinomycin combined with PIEZO1 stimulation. These treatments cancelled PIEZO1-induced *STC1* expression (Figure 3E). To further assess the specificity of HIF-1α involvement, we examined the hypoxia-inducible factor-2α (HIF-2α) inhibitor belzutifan, as HIF-2α activation is known to worsen PH *in vivo*.^22^ PIEZO1-induced *STC1* expression was unaffected by HIF-2α inhibition (Figure 3E), suggesting that hydrostatic pressure induces *STC1* expression via the PIEZO1–HIF-1α signaling axis.

### STC1 Inhibits PASMC Proliferation

To assess the effect of STC1 on PASMC proliferation, immunofluorescence staining for STC1, α-smooth muscle actin (α-SMA), and proliferating cell nuclear antigen (PCNA) was performed in lung sections from non-PAH controls and IPAH patients. In IPAH lungs, upregulated STC1 overlapped with α-SMA-positive and PCNA-positive regions, suggesting a role in proliferating PASMCs (Figure 4A).

**Figure 4.**
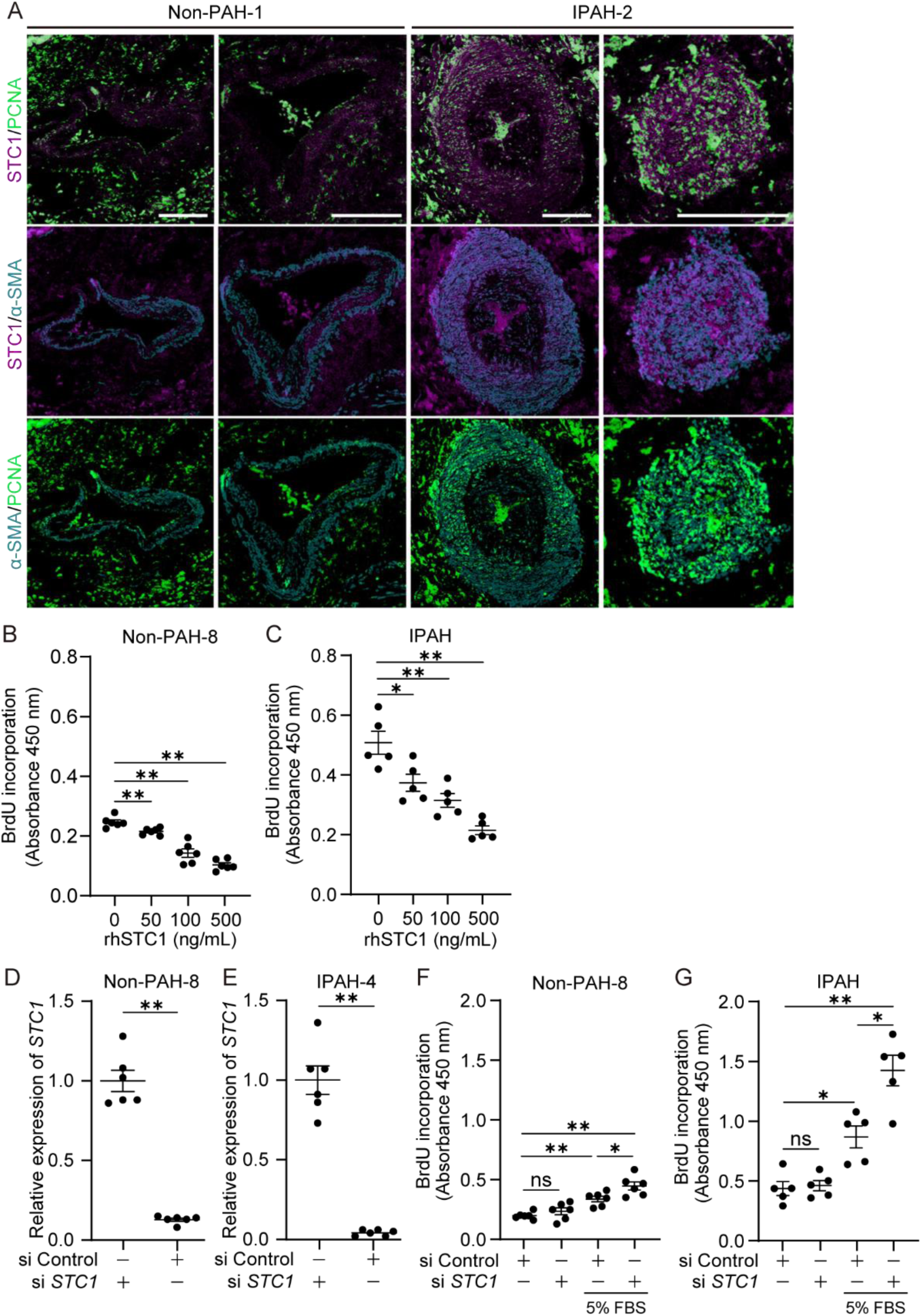
STC1 regulates cell proliferation in PASMCs. **A**, Representative images of immunofluorescence staining for stanniocalcin-1 (STC1, magenta), proliferating cell nuclear antigen (PCNA, green), and α-smooth muscle actin (α-SMA, cyan) in lung sections from a non-PAH control and a patient with idiopathic pulmonary arterial hypertension (IPAH). Scale bars: 50 μm. **B**-**C**, Effect of recombinant human STC1 (rhSTC1) on cell proliferation. Non-PAH-8 pulmonary artery smooth muscle cells (PASMCs; **B**) and IPAH-derived PASMCs (**C**). Proliferation was assessed by 5-bromo-2′-deoxyuridine (BrdU) incorporation assay and measured by ELISA. Donor-derived PASMCs were tested in a single lot with 6 replicates (n=6), and IPAH-derived PASMCs represent the mean value from each of 5 independent patient-derived PASMC line (n=5). Each dot represents an individual replicate (Non-PAH) or a patient-derived PASMCs line (IPAH). **D**-**E**, PASMCs were transfected with small interfering RNA (siRNA) targeting *STC1* (si *STC1*) or control siRNA (si Control), and knockdown efficiency was assessed by quantitative polymerase chain reaction (qPCR). Relative expression of *STC1* mRNA in a Non-PAH-8 PASMCs (n=6; **D**) and in an IPAH-4 PASMCs (n=6; **E**) is shown; the IPAH line selected was that with the highest basal expression of *STC1*. **F**-**G**, Proliferation following siRNA-mediated knockdown of *STC1*, measured by BrdU incorporation assay with ELISA readout in donor- and IPAH-derived PASMCs. Each dot represents data from an individual replicate (Non-PAH-8; **F**) or IPAH-derived PASMCs lines (**G**). Data are expressed as mean ± SEM. For comparisons among multiple groups, statistical significance was assessed using the Kruskal-Wallis test followed by post hoc Mann–Whitney U tests. Comparisons between two groups were performed using the Mann–Whitney U test. \**P* < 0.05, \*\**P* < 0.01, ns: not significant.

To evaluate the functional role of STC1, rhSTC1 was administered to PASMCs derived from non-PAH and IPAH patients. BrdU incorporation assays revealed a dose-dependent suppression of proliferation in both cell types following rhSTC1 treatment (Figures 4B-C), suggesting that STC1 negatively regulates PASMC proliferation. Furthermore, siRNA–mediated knockdown of *STC1* markedly increased BrdU incorporation (Figures 4D-G), indicating enhanced proliferative activity in the absence of STC1. Taken together, these results demonstrate that STC1 expression is upregulated in IPAH and acts as an endogenous negative regulator of PASMC proliferation.

### STC1 Increases Expression of Cell Cycle Arrest–Related Proteins

To elucidate the underlying molecular mechanisms of STC1-mediated inhibition of cell proliferation, rhSTC1 was administered to IPAH-derived PASMCs, and the expression of cell cycle arrest–related proteins was assessed by western blot analysis. First, we examined the time course of p38 phosphorylation, which is known to respond relatively early among cell cycle regulatory molecules.^23–25^ rhSTC1 stimulation promoted phosphorylation of p38, with the maximal response observed at 2 h after stimulation (Figure 5A), showing a significant increase (Figures 5B-C). Next, we analyzed the p53– p21 pathway, a reported downstream signal of p38,^26^ in samples collected 12 h after rhSTC1 administration. rhSTC1 stimulation increased the expression of phospho-p38, phospho-p53, p21, and p27, while reducing cyclin-dependent kinase 2 (CDK2) expression, indicating activation of cell cycle inhibitory pathways (Figures 5B-C).^27,28^

**Figure 5.**
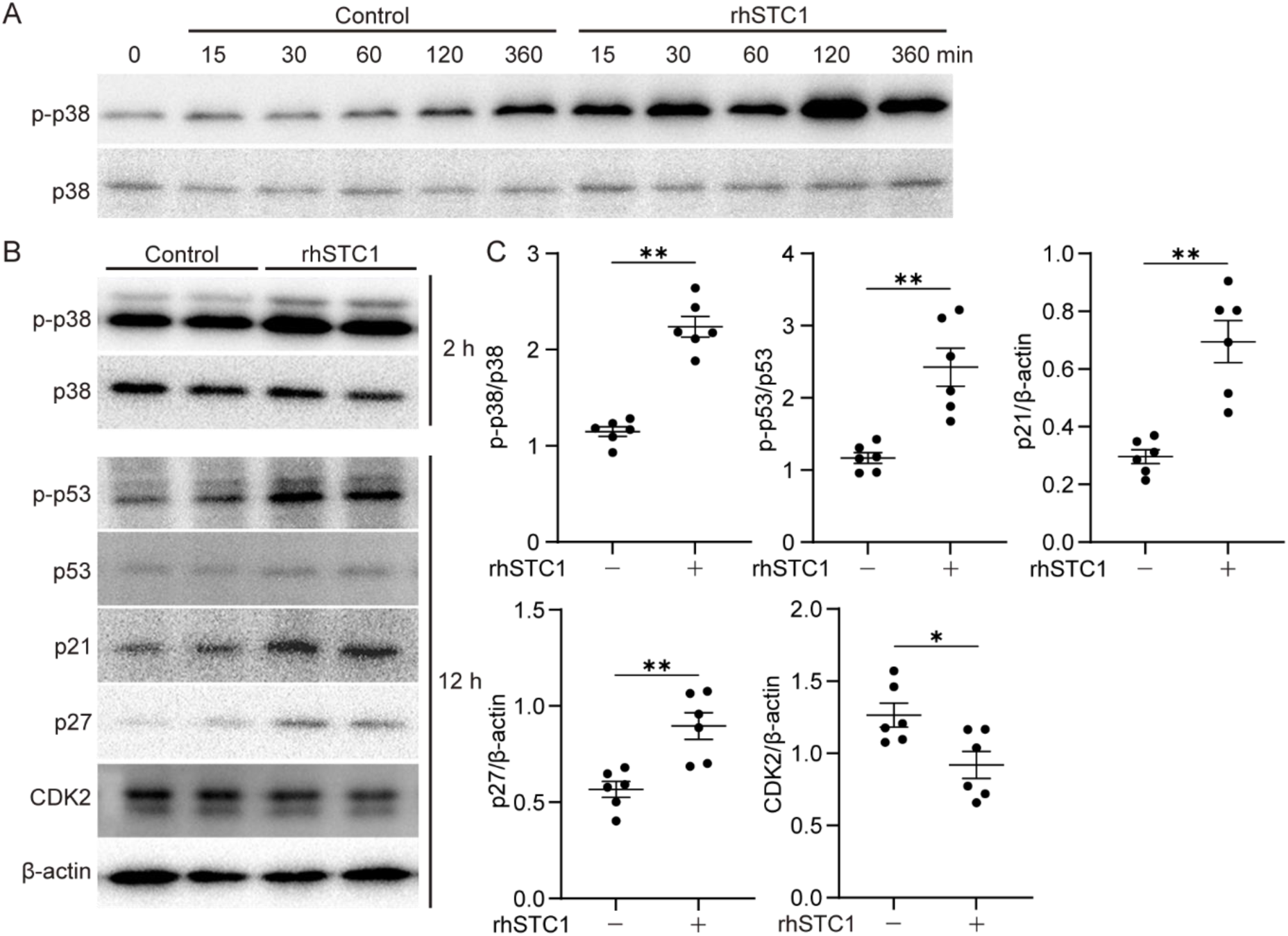
Effect of STC1 on cell cycle regulatory proteins in PASMCs. Pulmonary artery smooth muscle cells (PASMCs) derived from a patient with idiopathic pulmonary arterial hypertension (IPAH-4) were treated with recombinant human stanniocalcin-1 (rhSTC1, 100 ng/mL) and collected at predetermined time points for western blot analysis. **A**, Time course of phospho-p38 (p-p38) and total p38 expression at 0, 15, 30, 60, 120, and 360 min after rhSTC1 treatment. **B**, Representative western blot images showing expression of p-p38, total p38, phospho-p53 (p-p53), total p53, p21, p27, CDK2 and β-actin at the indicated time points (p38 and p-p38: 2 h; other proteins: 12 h). **C**, Quantitative analysis of western blot data. Levels of p-p38 were normalized to total p38, p-p53 to total p53, and other proteins to β-actin. Data are presented as mean ± SEM (n = 6). Statistical analysis was performed using the Mann–Whitney U test. \**P* < 0.05, \*\**P* < 0.01.

### Deficiency of STC1 Exacerbates Hypoxia-Induced Pulmonary Hypertension

To determine the *in vivo* role of STC1, we analyzed *Stc1*⁻^/^⁻ mice exposed to chronic hypoxia for 4 weeks (Figure 6A). Compared with WT controls, *Stc1*⁻^/^⁻ mice exhibited more severe PH, as indicated by increased RVSP (Figures 6B-C) and enhanced pulmonary arterial medial wall thickness (Figures 6D-E). Consistently, immunostaining revealed increased α-SMA-positive area under hypoxia, which was further augmented in *Stc1⁻^/^⁻* mice (Figure S3). Staining for von Willebrand factor, an endothelial marker, showed no overt change under conditions of hypoxia or *Stc1* deficiency (Figure S3). These differences occurred despite comparable age, body weight, and tibia length (Table S6). In this model, hypoxia increased Fulton index (RV/[LV+S]) in both genotypes; however, no significant difference was observed between WT and *Stc1*⁻^/^⁻ mice (Figure 6F). Systemic arterial pressure, left ventricular systolic pressure, and left ventricular ejection fraction were unaffected by either hypoxia or *Stc1* deficiency (Figures 6G-I). The same trends were observed when the data were analyzed separately in males and females (Figure S4).

**Figure 6.**
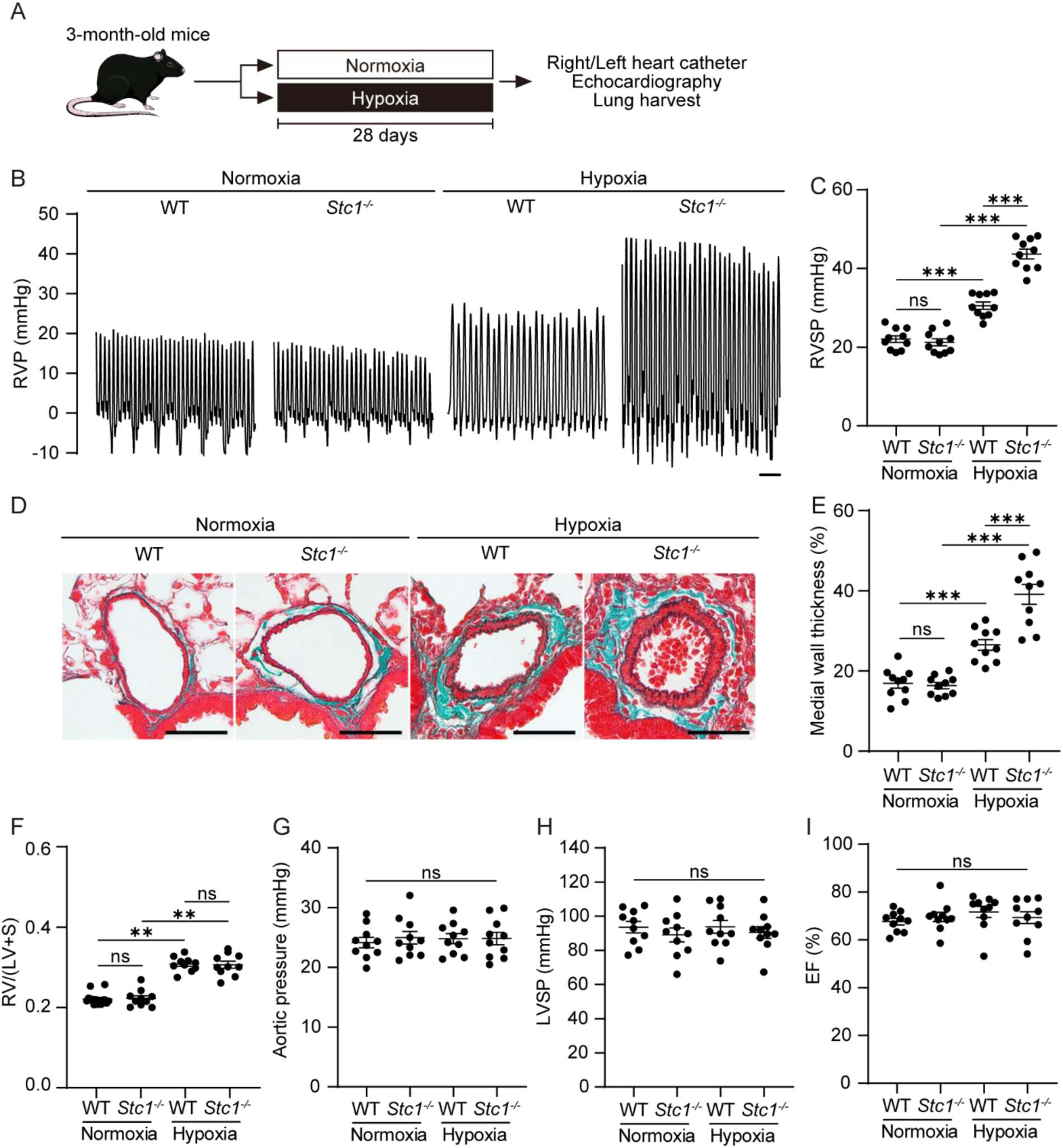
Loss of STC1 promotes the progression of hypoxia-induced pulmonary hypertension in mice. **A**, Experimental protocol for generating the hypoxia-induced pulmonary hypertension (PH) model. Wild-type (WT) and *Stc1*-knockout (*Stc1*⁻^/^⁻) mice were housed in normoxic (fraction of inspired oxygen [FiO_2_] 21%) or hypoxic (FiO_2_ 10%) chambers for 28 days, followed by physiological and histological assessments. **B**, Representative right ventricular pressure (RVP) waveforms from WT and *Stc1*⁻^/^⁻ mice under normoxic or hypoxic conditions. Scale bar: 5 s. **C**, Right ventricular systolic pressure (RVSP) in WT and *Stc1*⁻^/^⁻ mice under normoxic or hypoxic conditions (total n=10 per group; 5 males and 5 females). **D**, Representative lung sections stained with Elastica–Masson–Goldner. Scale bars: 50 μm. **E**, Quantification of pulmonary vascular remodeling, expressed as the percentage of medial wall thickness relative to total vessel diameter (total n=10 per group; 5 males and 5 females).Right ventricular hypertrophy index (calculated as RV/[LV+S], **F**), Aortic pressure (**G**), left ventricular systolic pressure (LVSP, **H**), and left ventricular ejection fraction (EF, **I**) in WT and *Stc1*⁻^/^⁻ mice exposed to normoxic or hypoxic conditions (total n=10 per group; 5 males and 5 females). Data are presented as mean ± SEM. Statistical analysis was performed using Kruskal–Wallis test followed by Mann–Whitney U test. \*\**P* < 0.01, \*\*\**P* < 0.001, ns: not significant.

To determine whether *Stc1* deficiency promotes PASMC proliferation *in vivo*, immunofluorescence staining for α-SMA and PCNA was performed on mouse lung sections. Under normoxia, no difference was observed between genotypes. However, under hypoxia, *Stc1*⁻^/^⁻ mice exhibited a significantly higher proportion of PCNA-positive PASMCs than WT controls (Figures 7A-B). We next evaluated perivascular macrophage infiltration, which has been reported to increase in pulmonary hypertension.^29,30^ CD68-positive macrophages, an M1 macrophage marker, were increased in *Stc1*⁻^/^⁻ mice even under normoxia and were further elevated under hypoxia (Figures 7C and 7D). In contrast, CD206-positive macrophages, an M2 macrophage marker, were increased under hypoxia in both genotypes without significant differences between groups (Figures 7E-F). Apoptosis was unaffected by either hypoxia or *Stc1* deficiency (Figure S5). Collectively, these findings demonstrate that loss of *Stc1* promotes PASMC proliferation and exacerbates the progression of hypoxia-induced pulmonary hypertension.

**Figure 7.**
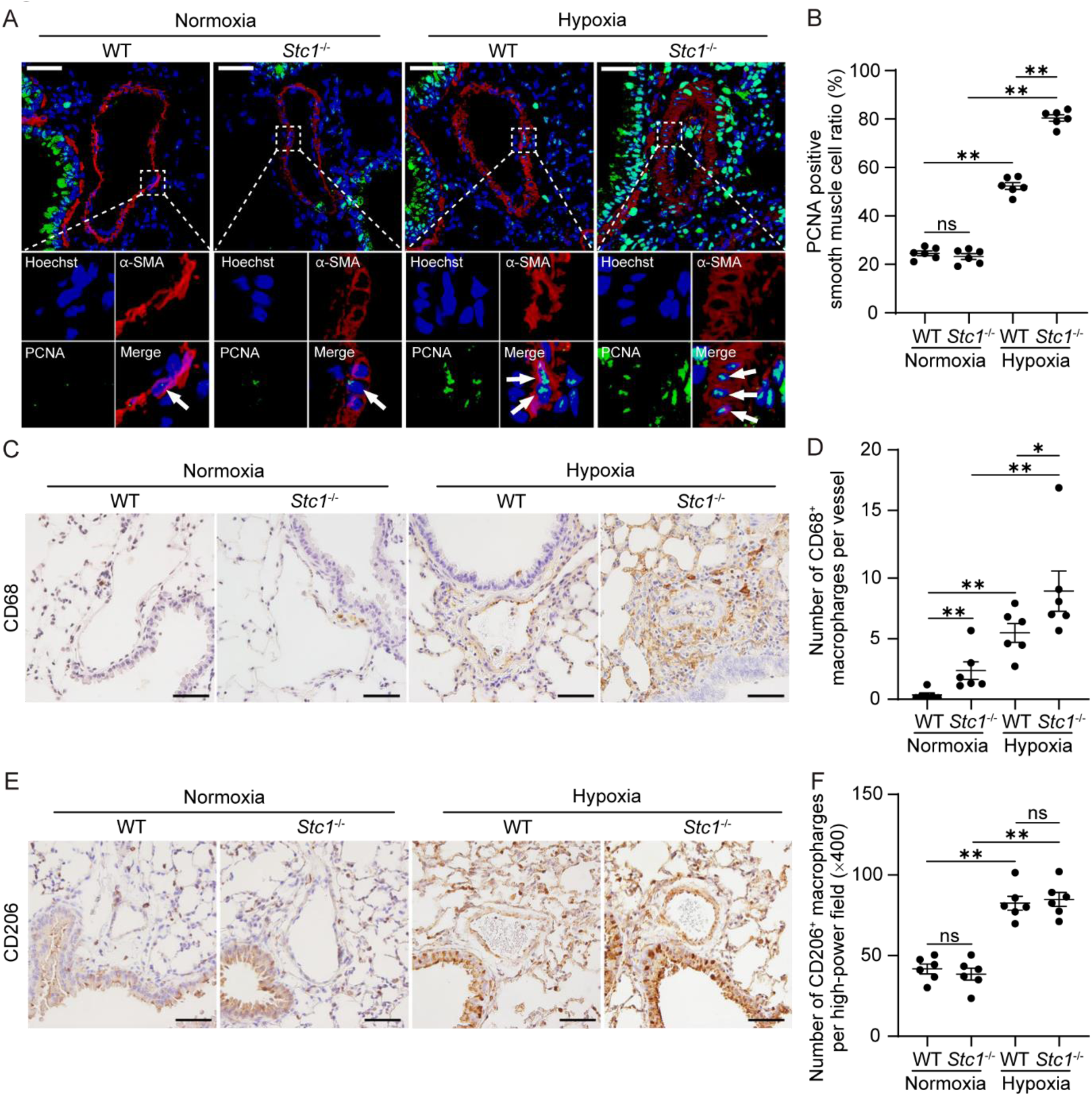
Proliferative and macrophage marker profiles in WT and Stc1⁻^/^⁻ mouse lungs under normoxic or hypoxic conditions. **A**, Representative immunofluorescence images of lung sections from wild-type (WT) and *Stc1*-knockout (*Stc1*⁻^/^⁻) mice stained for proliferating cell nuclear antigen (PCNA; green), α–smooth muscle actin (α-SMA; red), and Hoechst (blue) under normoxic or hypoxic conditions. Arrows indicate PCNA positive pulmonary artery smooth muscle cells (PASMCs). **B**, Quantification of PCNA-positive PASMCs as a percentage of total PASMCs in WT and *Stc1*⁻^/^⁻ mice under normoxic or hypoxic conditions (n=6 per group). **C**, Representative immunohistochemistry staining for CD68 in WT and Stc1⁻/⁻ mice exposed to normoxia or hypoxia. **D**, Quantification of CD68-positive cells per pulmonary artery. **E**, Representative immunohistochemistry staining for CD206 in WT and Stc1⁻/⁻ mice exposed to normoxia or hypoxia. **F**, Quantification of CD206-positive cells per high-power field. Data are presented as mean ± SEM. Statistical significance was determined by the Kruskal–Wallis test followed by the Mann–Whitney U test. \**P*<0.05, \*\**P*<0.01, ns: not significant. Scale bars: 50 μm.

### Recombinant STC1 Attenuates the Progression of Hypoxia-Induced Pulmonary Hypertension

To further evaluate the therapeutic potential of STC1 in pulmonary hypertension, rhSTC1 was administered via inhalation every 3 days throughout the hypoxic exposure period in a murine model of hypoxia-induced PH (Figure 8A). Repeated administration of rhSTC1 significantly reduced RVSP in both WT and *Stc1*⁻^/^⁻ mice compared with vehicle controls (Figures 8B-C). Consistent with the hemodynamic improvement, pulmonary arterial medial wall thickening was markedly attenuated in both genotypes following rhSTC1 treatment (Figures 8D-E), indicating a reduction in vascular remodeling.

**Figure 8.**
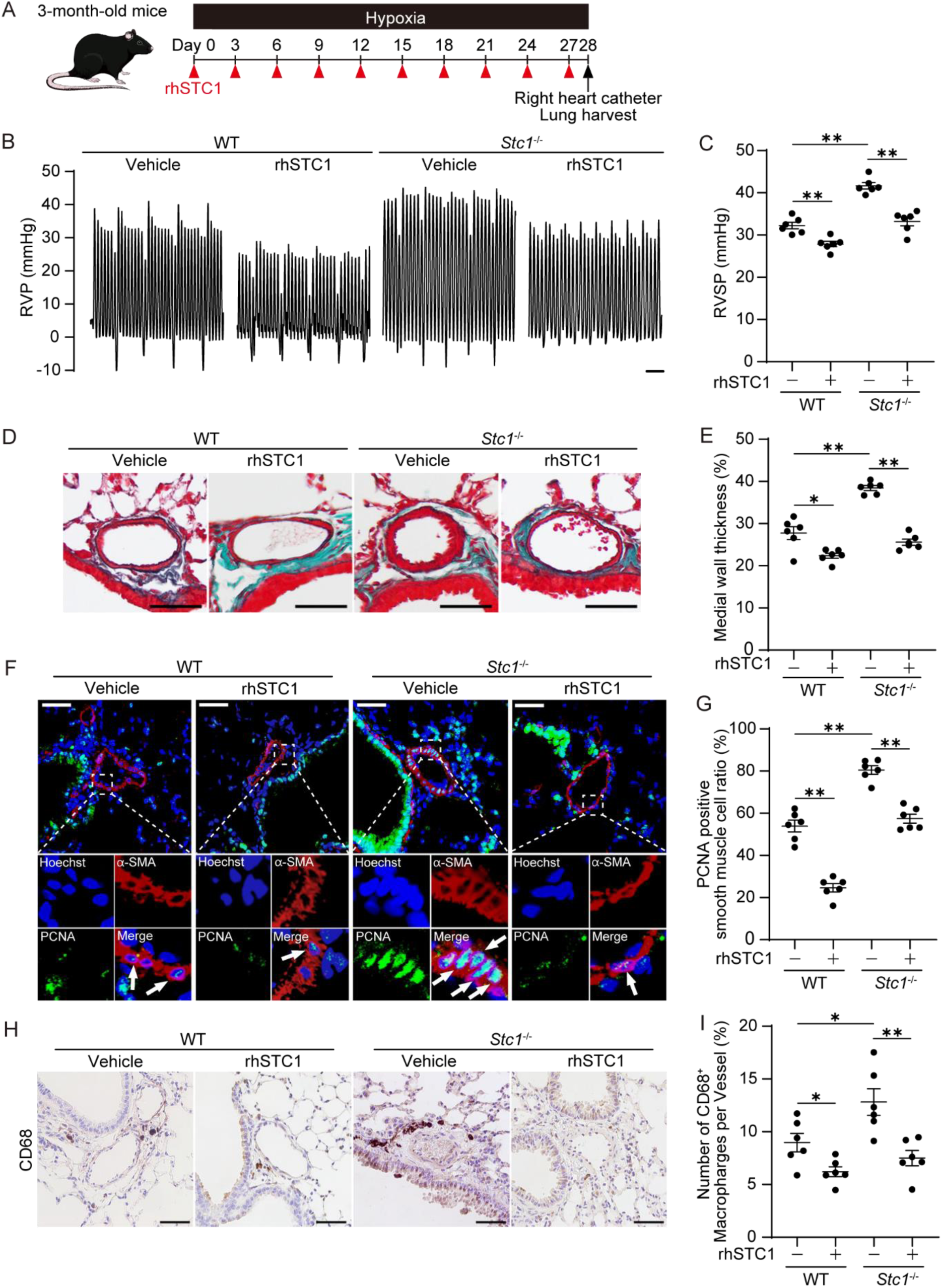
Recombinant STC1 attenuates the pathological features of hypoxia-induced pulmonary hypertension in mice. **A**, Experimental protocol for generating the hypoxia-induced pulmonary hypertension (PH) model mice and administering recombinant human STC1 (rhSTC1). Wild-type and *Stc1*⁻^/^⁻ mice were exposed to hypoxic conditions (FiO_2_ 10%) for 28 days. During this period, mice received either rhSTC1 (2 µg/ 50 µL PBS) or vehicle via inhalation every 3 days. Physiological and histological analyses were subsequently performed. **B**, Representative right ventricular pressure waveforms from WT and *Stc1*⁻^/^⁻ mice treated with rhSTC1 or vehicle. Scale bar: 5 s. **C**, Right ventricular systolic pressure (RVSP) in WT and *Stc1*⁻^/^⁻ mice treated with rhSTC1 or vehicle (n=6). **D**, Representative lung sections stained with Elastica–Masson–Goldner from WT and *Stc1*⁻^/^⁻ mice treated with rhSTC1 or vehicle. **E**, Quantification of pulmonary vascular remodeling by calculating the percentage of medial wall thickness relative to total vessel diameter in WT and *Stc1*⁻^/^⁻ mice treated with rhSTC1 or vehicle (n=6). **F**, Representative immunofluorescence staining of lung sections from WT and *Stc1*⁻^/^⁻ mice treated with rhSTC1 or vehicle for proliferating cell nuclear antigen (PCNA, green), α-smooth muscle actin (α-SMA, red), and Hoechst (blue). Arrows indicate PCNA positive pulmonary artery smooth muscle cells (PASMCs). **G**, Quantification of PCNA-positive PASMCs as a percentage of total PASMCs in WT and *Stc1*⁻^/^⁻ mice treated with rhSTC1 or vehicle (n=6). **H**, Representative immunohistochemistry staining for CD68 in WT and *Stc1*⁻^/^⁻ mice exposed to hypoxia. **I**, Quantification of CD68-positive cells per pulmonary artery. Data are presented as the mean ± SEM. Statistical analysis was performed using Kruskal– Wallis test followed by Mann–Whitney U test. \**P* < 0.05, \*\**P* < 0.01. Scale bars: 50 μm.

To investigate whether the observed structural improvement was accompanied by suppression of smooth muscle proliferation, immunofluorescence staining forα-SMA and PCNA was performed on lung sections obtained from hypoxia-exposed mice. The proportion of PCNA-positive PASMCs was significantly decreased in rhSTC1-treated groups in both WT and *Stc1*⁻^/^⁻ mice (Figures 8F-G), demonstrating that exogenous rhSTC1 effectively suppresses PASMC proliferative activity under hypoxic conditions.

The inflammatory component of pulmonary vascular remodeling was examined by assessing the perivascular infiltration of CD68-positive macrophages. rhSTC1 administration markedly reduced macrophage accumulation in both WT and *Stc1*⁻^/^⁻ lungs (Figures 8H-I), suggesting that STC1 exerts anti-inflammatory effects in addition to its antiproliferative action. Collectively, these findings demonstrate that exogenous rhSTC1 inhalation ameliorates the hypoxia-induced PH phenotype in *Stc1*⁻^/^⁻ mice and alleviates pulmonary vascular remodeling and right ventricular pressure overload in wild-type mice.

## Discussion

In the present study, we applied a custom-designed hydrostatic pressure apparatus capable of reproducing the pressure oscillations observed in the pulmonary arteries of patients with PAH. Transcriptomic analysis of IPAH-derived PASMCs subjected to this system revealed significant enrichment of multiple cell cycle-related gene sets by GSEA. These findings confirm that our apparatus is a platform that, at least in part, recapitulates the mechanical environment of PAH. Using this platform, we identified STC1 as a mechanosensitive molecule induced via PIEZO1–HIF-1α signaling, which exerted antiproliferative and protective effects in IPAH PASMCs. Furthermore, in a hypoxia-induced PAH model, *Stc1* deficiency aggravated disease severity, whereas supplementation ameliorated pathological remodeling, suggesting that STC1 represents a novel therapeutic target in PAH.

Previous studies have established PIEZO1 as a critical mechanosensor in vascular cells,^31^ capable of transducing mechanical stimuli into downstream signals such as Ca^2+^ influx and HIF-1α activation.^18,19^ However, the downstream effectors of PIEZO1 in PAH have remained largely undefined. In this study, we identified STC1 as a novel effector induced in a PIEZO1–HIF-1α–dependent manner in PASMCs derived from patients with IPAH. STC1 has been implicated in cell proliferation, oxidative stress regulation, and mitochondrial homeostasis in cancer cells and renal cells,^16,32^ but its role in VSMCs or in PH has not been previously elucidated. Importantly, the absence of STC1 induction by hydrostatic pressure in non-PAH PASMCs suggests that disease-specific contexts may amplify mechanosensitive transcriptional responses. In particular, the upregulation of PIEZO1 observed in IPAH^33,34^ may potentiate exaggerated responses to hydrostatic stress. Collectively, these findings extend our understanding of hydrostatic pressure– responsive signaling in PAH and establish STC1 as a previously unrecognized protective mediator.

The finding that rhSTC1 induced cell cycle arrest and suppressed pulmonary vascular remodeling supports the concept that increased STC1 may function as a protective mechanism against pathological remodeling. In human lung tissues, STC1 expression was elevated in IPAH compared with donors; however, the proliferation marker PCNA was also increased. This suggests a state of relative STC1 insufficiency in IPAH, resembling the phenomenon in which B-type natriuretic peptide (BNP) is upregulated in response to hemodynamic stress and exerts cardioprotective effects.^35^ Indeed, circulating STC1 levels have been recently reported to predict adverse outcomes in patients with IPAH.^36^ Taken together, these findings indicate that, similar to BNP, STC1 is increased as a reflection of disease progression but exerts protective effects against pathological remodeling, and that exogenous supplementation of STC1 may represent a novel therapeutic strategy for PAH.

STC1 suppressed PASMC proliferation and attenuated PAH progression *in vivo*, indicating its protective role in pulmonary vascular remodeling. Therapeutic enhancement of STC1 signaling could be achieved either by supplementation of STC1 or by upstream modulation of PIEZO1-HIF-1α activity. However, PIEZO1 and HIF-1α pathways exert pleiotropic functions *in vivo*,^37–39^ particularly in the vasculature, where they promote remodeling to maintain oxygen delivery and vascular tone.^39^ Thus, supplementation of STC1, a downstream effector, may represent a safer and more specific strategy. Only a limited number of receptors for STC1 have been described. insulin-like growth factor 2 receptor (IGF2R) has been suggested as a binding partner, but it primarily functions in insulin-like growth factor 2 (IGF2) clearance and transforming growth factor β (TGF-β) regulation and is not directly implicated in the control of vascular tone.^40^ Therefore, STC1 is more likely to target proliferative and remodeling processes in PAH, potentially complementing existing vasodilator therapies.

IL-6, a central inflammatory cytokine in PAH,^41^ is produced by M1 macrophages, which have been reported to be increased in PAH.^29,30^ STC1 has been shown to exert inhibitory effects on IL-6 upregulation^42^ and, in the present study, STC1 suppressed perivascular infiltration of M1 macrophages. Furthermore, STC1 has been reported to suppress fibrosis induced by TGF-β, a major driver of vascular remodeling.^32^ Taken together, these findings suggest that STC1 functions not only as an antiproliferative factor in PASMCs but also as a regulator of anti-inflammatory and antifibrotic pathways, thereby highlighting its multifaceted therapeutic potential in PAH.

Several limitations warrant consideration. First, although we demonstrated that PIEZO1–HIF-1α signaling mediates STC1 induction, the full transcriptional network upstream of STC1 remains to be elucidated. Second, the number of donor-derived PASMC lines was small, and inter-patient variability may influence mechanosensitive transcriptional responses. Third, the hypoxia-induced PH model captures many aspects of PAH but does not fully replicate the complex hemodynamic and molecular environment of human IPAH. Finally, mechanosensitive channels other than PIEZO1 or integrin-mediated pathways may contribute to STC1 induction.

In conclusion, we identified the PIEZO1–HIF-1α–STC1 axis as a novel mechanosensitive pathway in IPAH PASMCs. Using a custom hydrostatic pressure apparatus that replicates disease-relevant mechanical stress, we demonstrated that STC1 is upregulated in response to pathological pressure and exerts antiproliferative, anti-inflammatory, and protective effects against pulmonary vascular remodeling. These findings provide new mechanistic insight into how abnormal hemodynamic forces contribute to PAH and highlight STC1 as a promising therapeutic target.

## Acknowledgements

We thank Yuko Hidaka, Yuka Sawada and Tomoko Furuta (Department of Physiology, Tokyo Medical University) for their technical assistance. We are also deeply grateful to Drs. Hiromu Tanaka, Hideto Iizuka, and Koichi Fukunaga (Department of Respiratory Medicine, Keio University School of Medicine) for providing expert instruction on intratracheal drug administration in mice. The generation of the custom-built periodic hydrostatic pressure culture system was aided by Koganei Co., Ltd (Tokyo, Japan).

## Source of Funding

This work was supported by MEXT/JSPS KAKENHI (MK, JP23K07637), and a research grant from Nippon Shinyaku Research Grant (2024), and partly supported by MEXT/JSPS KAKENHI (YK, JP21K07352; KN, JP21K08108; UY, JP24K02427, JP23K18320), and the Japan Agency for Medical Research and Development (AMED) (UY, 23ek0210183, 24ek0109773).

## Disclosure

All authors report no conflicts of interest.

## Supplemental Materials

Supplemental Methods

Tables S1-S6

Figures S1-S5

## Non-standard Abbreviations and Acronyms

ACTA2: smooth muscle α-actin
AP-1: activator protein-1
α-SMA: alpha-smooth muscle actin
BNP: B-type natriuretic peptide
BrdU: bromodeoxyuridine
CDK2: cyclin-dependent kinase 2
FOSL1: FOS like 1, AP-1 transcription factor subunit
GOBP: Gene Ontology Biological Process
GSEA: Gene Set Enrichment Analysis
HIF-1α: hypoxia-inducible factor-1 alpha
HIF-2α: hypoxia-inducible factor-2 alpha
IGF2: insulin-like growth factor 2
IGF2R: insulin-like growth factor 2 receptor
IPAH: idiopathic pulmonary arterial hypertension
KEGG: Kyoto Encyclopedia of Genes and Genomes
KLF4: Krüppel-like factor 4
MYH11: myosin heavy chain 11
NES: normalized enrichment score
PASMC: pulmonary arterial smooth muscle cell
PAH: pulmonary arterial hypertension
PCNA: proliferating cell nuclear antigen
PH: pulmonary hypertension
PIEZO1: piezo type mechanosensitive ion channel component 1
rhSTC1: recombinant human stanniocalcin-1
RVP: right ventricular pressure
RVSP: right ventricular systolic pressure
siRNA: small interfering RNA
STC1: stanniocalcin-1
TAGLN: transgelin
TGF-β: transforming growth factor beta
TRPM7: transient receptor potential melastatin 7
TRPV4: transient receptor potential vanilloid 4
TUNEL: terminal deoxynucleotidyl transferase dUTP nick end labeling
VSMC: vascular smooth muscle cell
WT: wild-type

## Author Contributions

- Study conception and design: Utako Yokoyama, Mariko Kogami
- Data acquisition: Mariko Kogami, Yuko Kato, Satoko Ito, Hana Inoue, Shota Tanifuji, Mayumi Yokotsuka, Yoshinari Yamamoto
- Analysis and interpretation: Mariko Kogami, Utako Yokoyama, Keiko Uchida, Toshitaka Nagao, Shinji Abe, Roger R. Reddel, Kazufumi Nakamura
- Drafting of manuscript: Mariko Kogami, Utako Yokoyama
- Critical revision: Utako Yokoyama
- Final approval of the version to be published: All authors

## Notes

### Competing Interest Statement

The authors have declared no competing interest.

## References

1. Humbert M, Kovacs G, Hoeper MM, Badagliacca R, Berger RMF, Brida M, Carlsen J, Coats AJS, Escribano-Subias P, Ferrari P, et al. 2022 ESC/ERS Guidelines for the diagnosis and treatment of pulmonary hypertension. Eur Heart J. 2022;43:3618–3731. doi: 10.1093/eurheartj/ehac237

2. Lechartier B, Berrebeh N, Huertas A, Humbert M, Guignabert C, Tu L. Phenotypic Diversity of Vascular Smooth Muscle Cells in Pulmonary Arterial Hypertension: Implications for Therapy. Chest. 2022;161:219–231. doi: 10.1016/j.chest.2021.08.040

3. Wang A, Valdez-Jasso D. Cellular mechanosignaling in pulmonary arterial hypertension. Biophys Rev. 2021;13:747–756. doi: 10.1007/s12551-021-00828-3

4. Prystopiuk V, Fels B, Simon CS, Liashkovich I, Pasrednik D, Kronlage C, Wedlich-Söldner R, Oberleithner H, Fels J. A two-phase response of endothelial cells to hydrostatic pressure. J Cell Sci. 2018;131. doi: 10.1242/jcs.206920

5. Yu Y, Cai Y, Yang F, Yang Y, Cui Z, Shi D, Bai R. Vascular smooth muscle cell phenotypic switching in atherosclerosis. Heliyon. 2024;10:e37727. doi: 10.1016/j.heliyon.2024.e37727

6. Cao G, Xuan X, Hu J, Zhang R, Jin H, Dong H. How vascular smooth muscle cell phenotype switching contributes to vascular disease. Cell Commun Signal. 2022;20:180. doi: 10.1186/s12964-022-00993-2

7. Kojima T, Nakamura T, Saito J, Hidaka Y, Akimoto T, Inoue H, Chick CN, Usuki T, Kaneko M, Miyagi E, et al. Hydrostatic pressure under hypoxia facilitates fabrication of tissue-engineered vascular grafts derived from human vascular smooth muscle cells *in vitro*. Acta Biomater. 2023;171:209–222. doi: 10.1016/j.actbio.2023.09.041

8. Yokoyama U, Tonooka Y, Koretake R, Akimoto T, Gonda Y, Saito J, Umemura M, Fujita T, Sakuma S, Arai F, et al. Arterial graft with elastic layer structure grown from cells. Sci Rep. 2017;7:140. doi: 10.1038/s41598-017-00237-1

9. Chang AC, Cha J, Koentgen F, Reddel RR. The murine stanniocalcin 1 gene is not essential for growth and development. Mol Cell Biol. 2005;25:10604–10610. doi: 10.1128/mcb.25.23.10604-10610.2005

10. Ono M, Ohkouchi S, Kanehira M, Tode N, Kobayashi M, Ebina M, Nukiwa T, Irokawa T, Ogawa H, Akaike T, et al. Mesenchymal stem cells correct inappropriate epithelial-mesenchyme relation in pulmonary fibrosis using stanniocalcin-1. Mol Ther. 2015;23:549–560. doi: 10.1038/mt.2014.217

11. Kato Y, Yokoyama U, Yanai C, Ishige R, Kurotaki D, Umemura M, Fujita T, Kubota T, Okumura S, Sata M, et al. Epac1 Deficiency Attenuated Vascular Smooth Muscle Cell Migration and Neointimal Formation. Arterioscler Thromb Vasc Biol. 2015;35:2617–2625. doi: 10.1161/atvbaha.115.306534

12. Okumura S, Oka S, Sasaki T, Cooley MA, Hidaka Y, Inoue H, Nishijima H, Ohno SI, Tanifuji S, Kaneko M, et al. Spatiotemporal EP4-fibulin-1 expression is associated with vascular intimal hyperplasia. Cardiovasc Res. 2024;120:2293–2306. doi: 10.1093/cvr/cvae211

13. Takahashi L, Ishigami T, Tomiyama H, Kato Y, Kikuchi H, Tasaki K, Yamashita J, Inoue S, Taguri M, Nagao T, et al. Increased Plasma Levels of Myosin Heavy Chain 11 Is Associated with Atherosclerosis. J Clin Med. 2021;10. doi: 10.3390/jcm10143155

14. Yap C, Mieremet A, de Vries CJM, Micha D, de Waard V. Six Shades of Vascular Smooth Muscle Cells Illuminated by KLF4 (Krüppel-Like Factor 4). Arterioscler Thromb Vasc Biol. 2021;41:2693–2707. doi: 10.1161/atvbaha.121.316600

15. Sobolev VV, Khashukoeva AZ, Evina OE, Geppe NA, Chebysheva SN, Korsunskaya IM, Tchepourina E, Mezentsev A. Role of the Transcription Factor FOSL1 in Organ Development and Tumorigenesis. Int J Mol Sci. 2022;23. doi: 10.3390/ijms23031521

16. Guo F, Li Y, Wang J, Li Y, Li Y, Li G. Stanniocalcin1 (STC1) Inhibits Cell Proliferation and Invasion of Cervical Cancer Cells. PLoS One. 2013;8:e53989. doi: 10.1371/journal.pone.0053989

17. Costa BP, Schein V, Zhao R, Santos AS, Kliemann LM, Nunes FB, Cardoso JCR, Félix RC, Canário AVM, Brum IS, et al. Stanniocalcin-1 protein expression profile and mechanisms in proliferation and cell death pathways in prostate cancer. Mol Cell Endocrinol. 2020;502:110659. doi: 10.1016/j.mce.2019.110659

18. Wang Z, Chen J, Babicheva A, Jain PP, Rodriguez M, Ayon RJ, Ravellette KS, Wu L, Balistrieri F, Tang H, et al. Endothelial upregulation of mechanosensitive channel Piezo1 in pulmonary hypertension. Am J Physiol Cell Physiol. 2021;321:C1010–c1027. doi: 10.1152/ajpcell.00147.2021

19. Chen Y, Yin Y, Luo M, Wu J, Chen A, Deng L, Xie L, Han X. Occlusal Force Maintains Alveolar Bone Homeostasis via Type H Angiogenesis. J Dent Res. 2023;102:1356–1365. doi: 10.1177/00220345231191745

20. Ye X, Xia Y, Zheng Y, Chen W, Chen Z, Cheng Z, Wang B. The function of Piezo1 in hepatoblastoma metastasis and its potential transduction mechanism. Heliyon. 2022;8:e10301. doi: 10.1016/j.heliyon.2022.e10301

21. Wang X, Cheng G, Miao Y, Qiu F, Bai L, Gao Z, Huang Y, Dong L, Niu X, Wang X, et al. Piezo type mechanosensitive ion channel component 1 facilitates gastric cancer omentum metastasis. J Cell Mol Med. 2021;25:2238–2253. doi: 10.1111/jcmm.16217

22. Shimoda LA. What’s HIF Got to Do with It? HIF-2 Inhibition and Pulmonary Hypertension. Am J Respir Crit Care Med. 2018;198:1363–1365. doi: 10.1164/rccm.201806-1130ED

23. Thornton TM, Rincon M. Non-classical p38 map kinase functions: cell cycle checkpoints and survival. Int J Biol Sci. 2009;5:44–51. doi: 10.7150/ijbs.5.44

24. Whitaker RH, Cook JG. Stress Relief Techniques: p38 MAPK Determines the Balance of Cell Cycle and Apoptosis Pathways. Biomolecules. 2021;11. doi: 10.3390/biom11101444

25. García-Flores N, Jiménez-Suárez J, Garnés-García C, Fernández-Aroca DM, Sabater S, Andrés I, Fernández-Aramburo A, Ruiz-Hidalgo MJ, Belandia B, Sanchez-Prieto R, et al. P38 MAPK and Radiotherapy: Foes or Friends? Cancers (Basel). 2023;15. doi: 10.3390/cancers15030861

26. Suhail TV, Singh P, Manna TK. Suppression of centrosome protein TACC3 induces G1 arrest and cell death through activation of p38-p53-p21 stress signaling pathway. Eur J Cell Biol. 2015;94:90–100. doi: 10.1016/j.ejcb.2014.12.001

27. García-Osta A, Dong J, Moreno-Aliaga MJ, Ramirez MJ. p27, The Cell Cycle and Alzheimeŕs Disease. Int J Mol Sci. 2022;23. doi: 10.3390/ijms23031211

28. Engeland K. Cell cycle regulation: p53-p21-RB signaling. Cell Death Differ. 2022;29:946–960. doi: 10.1038/s41418-022-00988-z

29. Hemnes AR, Luther JM, Rhodes CJ, Burgess JP, Carlson J, Fan R, Fessel JP, Fortune N, Gerszten RE, Halliday SJ, et al. Human PAH is characterized by a pattern of lipid-related insulin resistance. JCI Insight. 2019;4. doi: 10.1172/jci.insight.123611

30. Pugliese SC, Kumar S, Janssen WJ, Graham BB, Frid MG, Riddle SR, El Kasmi KC, Stenmark KR. A Time- and Compartment-Specific Activation of Lung Macrophages in Hypoxic Pulmonary Hypertension. J Immunol. 2017;198:4802–4812. doi: 10.4049/jimmunol.1601692

31. Davis MJ, Earley S, Li YS, Chien S. Vascular mechanotransduction. Physiol Rev. 2023;103:1247–1421. doi: 10.1152/physrev.00053.2021

32. Yang EM, Park JS, Joo SY, Bae EH, Ma SK, Kim SW. Stanniocalcin-1 suppresses TGF-β-induced mitochondrial dysfunction and cellular fibrosis in human renal proximal tubular cells. Int J Mol Med. 2022;50. doi: 10.3892/ijmm.2022.5163

33. Knoepp F, Abid S, Houssaini A, Lipskaia L, Gökyildirim MY, Born E, Marcos E, Arhatte M, Glogowska E, Vienney N, et al. Piezo1 in PASMCs: Critical for Hypoxia-Induced Pulmonary Hypertension Development. Circ Res. 2025;136:1031–1048. doi: 10.1161/circresaha.124.325475

34. Liao J, Lu W, Chen Y, Duan X, Zhang C, Luo X, Lin Z, Chen J, Liu S, Yan H, et al. Upregulation of Piezo1 (Piezo Type Mechanosensitive Ion Channel Component 1) Enhances the Intracellular Free Calcium in Pulmonary Arterial Smooth Muscle Cells From Idiopathic Pulmonary Arterial Hypertension Patients. Hypertension. 2021;77:1974–1989. doi: 10.1161/hypertensionaha.120.16629

35. Kuwahara K. The natriuretic peptide system in heart failure: Diagnostic and therapeutic implications. Pharmacol Ther. 2021;227:107863. doi: 10.1016/j.pharmthera.2021.107863

36. Yang Y, Shu SR, Sun B, Wang PZ, Zeng QX, Wang CS, Xiong CM. Improved prognosis prediction of idiopathic pulmonary arterial hypertension using targeted plasma proteomics in disease management. European Heart Journal. 2024;45:ehae666.2241. doi: 10.1093/eurheartj/ehae666.2241

37. Zhang Y, Zou W, Dou W, Luo H, Ouyang X. Pleiotropic physiological functions of Piezo1 in human body and its effect on malignant behavior of tumors. Front Physiol. 2024;15:1377329. doi: 10.3389/fphys.2024.1377329

38. Huang JQ, Zhang H, Guo XW, Lu Y, Wang SN, Cheng B, Dong SH, Lyu XL, Li FS, Li YW. Mechanically Activated Calcium Channel PIEZO1 Modulates Radiation-Induced Epithelial-Mesenchymal Transition by Forming a Positive Feedback With TGF-β1. Front Mol Biosci. 2021;8:725275. doi: 10.3389/fmolb.2021.725275

39. Shinge SAU, Zhang D, Achu Muluh T, Nie Y, Yu F. Mechanosensitive Piezo1 Channel Evoked-Mechanical Signals in Atherosclerosis. J Inflamm Res. 2021;14:3621–3636. doi: 10.2147/jir.S319789

40. Wan HT, Ng AH, Lee WK, Shi F, Wong CK. Identification and characterization of a membrane receptor that binds to human STC1. Life Sci Alliance. 2022;5. doi: 10.26508/lsa.202201497

41. Xu WJ, Wu Q, He WN, Wang S, Zhao YL, Huang JX, Yan XS, Jiang R. Interleukin-6 and pulmonary hypertension: from physiopathology to therapy. Front Immunol. 2023;14:1181987. doi: 10.3389/fimmu.2023.1181987

42. Yamamoto K, Tajima Y, Hasegawa A, Takahashi Y, Kojima M, Watanabe R, Sato K, Shichiri M, Watanabe T. Contrasting effects of stanniocalcin-related polypeptides on macrophage foam cell formation and vascular smooth muscle cell migration. Peptides. 2016;82:120–127. doi: 10.1016/j.peptides.2016.06.009

43. Ono M, Ohkouchi S, Kanehira M, Tode N, Kobayashi M, Ebina M, Nukiwa T, Irokawa T, Ogawa H, Akaike T, Okada Y, Kurosawa H, Kikuchi T and Ichinose M. Mesenchymal stem cells correct inappropriate epithelial-mesenchyme relation in pulmonary fibrosis using stanniocalcin-1. Mol Ther. 2015;23:549–60.

44. Okumura S, Oka S, Sasaki T, Cooley MA, Hidaka Y, Inoue H, Nishijima H, Ohno SI, Tanifuji S, Kaneko M, Abe T, Kuroda M, Yokosuka T, Breyer RM, Homma H, Kato Y and Yokoyama U. Spatiotemporal EP4-fibulin-1 expression is associated with vascular intimal hyperplasia. Cardiovasc Res. 2024;120:2293–2306.

